# Phased diploid genome assemblies for three strains of *Candida albicans* from oak trees

**DOI:** 10.1101/697524

**Authors:** Jennafer A. P. Hamlin, Guilherme Dias, Casey M. Bergman, Douda Bensasson

**Author notes:** Equally contributing authors (Jennafer A. P. Hamlin, Guilherme Dias). Corresponding author: Douda Bensasson, Department of Plant Biology, University of Georgia, Miller Plant Sciences, Athens, GA 30602.

## Abstract

Although normally a harmless commensal, *Candida albicans* has the potential to generate a wide range of infections including systemic candidaemia, making it the most common cause of bloodstream infections worldwide with a high rate of mortality. *C. albicans* has long been considered an obligate commensal, however, recent studies suggest it can live outside animal hosts. Here, we have generated PacBio sequencing and phased genome assemblies for three *C. albicans* strains from oak trees in the United Kingdom (NCYC 4144, NCYC 4145, and NCYC 4146). Our results provide phased *de novo* diploid assemblies for *C. albicans* and provide a framework to study patterns of genomic variation within and among strains of an important fungal pathogen.

## INTRODUCTION

The fungus *Candida albicans* is found in the healthy human gut flora as a lifelong, harmless commensal (Odds *et al.* 2007; Barnett 2008). However, under certain circumstances, *C. albicans* can cause human yeast infections ranging from superficial infections of the skin to life-threatening systemic diseases or candidaemia (Odds *et al.* 2007; Magill *et al.* 2014). While typically considered an obligate commensal, recent work has shown that *C. albicans* can be isolated from oak trees in the UK (Bensasson *et al.* 2019) and fruits, soil, and plant matter in the USA (Opulente *et al.* 2019), providing evidence that *C. albicans* can also live outside animal hosts. *C. albicans* strains from oak trees are closely related to pathogenic strains, and comparison of strains from these different environments may be useful for identifying the genetic determinants of pathogenesis (Bensasson *et al.* 2019).

Genetic and genomic studies have confirmed that the *C. albicans* genome typically exists in a diploid heterozygous state (Barnett 2008; Hickman *et al.* 2013; Hirakawa *et al.* 2015; Ropars *et al.* 2018). The heterozygous nature of the *C. albicans* genome makes it particularly hard to assemble with current methods that assume a haploid or highly inbred genome (Jones *et al.* 2004; Nantel 2006). The initial effort to assemble the genome of the diploid *C. albicans* reference strain SC5314 performed a haploid assembly of Sanger shotgun sequences followed by an effort to separate (or “phase”) haplotypes in the subset of the genome where homologous chromosomes had sufficient divergence to be assembled as distinct contigs (Jones *et al.* 2004). Subsequently, van het Hoog *et al.* (2007) integrated contigs from Jones *et al.* (2004) with optical maps and other data to generate a haploid representation of all eight *C. albicans* chromosomes. Muzzey *et al.* (2013) then mapped short read sequences from SC5314 and a panel of diploid strains with loss of heterozygosity (LOH) for particular chromosomes (Legrand *et al.* 2008) to the haploid assembly from van het Hoog *et al.* (2007) to phase SNPs and short indel variants that distinguish the two haplotypes in SC5314. More recently, long-read sequencing has been used to generate a *de novo* haploid assembly of a pathogenic strain of *C. albicans* (Panthee *et al.* 2018). However, no phased *de novo* diploid genome assembly has currently been reported for pathogenic or environmental strains of *C. albicans*.

Phasing haplotypes is important for understanding the relationship between DNA sequence and phenotype in diploid organisms (reviewed in Tewhey *et al.* 2011). Phased genomes can also expose misassemblies in reference genomes (Korlach *et al.* 2017), identify point mutations and large structural variation between haplotypes (Zhou *et al.* 2019b), and allow detection of allele-specific expression and epigenetic modifications (Zhou *et al.* 2019a). In a predominantly asexual species such as *C. albicans* (Birky 1996; Barnett 2008; Bougnoux *et al.* 2008; Hirakawa *et al.* 2015), haplotype phasing of alleles is needed for correct inference of phylogenetic relationships (Birky 1996), and for detecting rare mating events that have potential implications for understanding adaptation to antifungal drugs (Berman and Hadany 2012) and disease emergence (Ropars *et al.* 2018).

Despite their utility, phased diploid representations of genomes are not commonly obtained during *de novo* genome assembly since phasing parental haplotypes is technically challenging. As of May 2019, there were only ~120 assemblies with some level of haplotype resolution in the NCBI Assembly database, a small number compared to well over 300,000 haploid assemblies (Kitts *et al.* 2016). Since for many organisms it is often not feasible or practical to construct strains with single haplotypes to circumvent the challenges of diploid assembly, different computational strategies have been developed to directly assemble diploid genomes. One such method is called FALCON-Unzip, which involves using heterozygous variation captured by PacBio Single-Molecule RealTime (SMRT) long-read sequencing to generate a *de novo* diploid assembly with haplotypes phased in heterozygous regions (Chin *et al.* 2016).

Here, we generated whole genome shotgun sequences using the PacBio SMRT technology (Eid *et al.* 2009) and applied the FALCON/FALCON-Unzip (Chin *et al.* 2016) assembly and phasing pipeline to three strains of *C. albicans* from oak trees in the UK (Bensasson *et al.* 2019). Our results demonstrate the feasibility of generating essentially-complete *de novo* diploid assemblies of *C. albicans* genomes using long-read sequencing and provide resources for future analysis of genome evolution and function in this important human fungal pathogen.

## MATERIALS AND METHODS

### Yeast strains

DNA was extracted from three strains of *C. albicans*, NCYC 4144 (FRI10b.1), NCYC 4145 (FRI11a.1) and NCYC 4146 (FRI5d.SM), which were isolated from oak tree bark (Robinson *et al.* 2016). The strains analyzed here were previously characterized using Illumina short-read sequencing, are all phylogenetically distinct and show unusually high heterozygosity (Bensasson *et al.* 2019). More specifically, comparison between the genome-wide SNPs of these oak strains and representatives from the known *C. albicans* clades and singletons described in Ropars *et al.* (2018) showed that NCYC 4146 belongs to clade 4, NCYC 4144 to clade 18 and NCYC 4145 does not resemble any known genome (Bensasson *et al.* 2019).

### Extraction of high molecular weight genomic DNA

For extraction of high molecular weight DNA, we used the Promega Wizard® Genomic DNA Purification Kit (A1125), and modified it to include an extra RNA digestion step, ethanol precipitation, longer centrifugation steps at higher centrifugal force and more reactions per sample (general recommendations are detailed in Bensasson 2018). For each strain, a single colony was used to inoculate 15 ml of YPD media and grown for 20-24 hours at 30°C. Cells were pelleted in two minutes at a relative centrifugal force of 16,162 g in 1.4 ml volumes in thirteen (NCYC 4145), fourteen (NCYC 4146) or eleven (NCYC 4144) 1.5 ml tubes. Cell walls were digested overnight at 37°C using 100 units lyticase (Sigma, L2524-50KU). Cells in each tube were pelleted (2 minutes at 16,128 g), resuspended in 300 µl Promega Nuclei Lysis solution and 100 µl Promega Protein Precipitation solution, cooled on ice for 5-30 minutes and cell debris was pelleted at 16,162 g for 10 minutes. The DNA in the supernatant was precipitated in 300 µl isopropanol and pelleted at 16,162 g for 10 minutes, washed with 300 µl of 70% ethanol at room temperature, pelleted at 16,162 g for 5 minutes, and air dried for 15 minutes. Pellets were resuspended in Promega DNA rehydration solution (1x TE buffer). RNA was digested using 1.5 µl (5.25 units) Promega RNase solution at 37°C for 1-2 hours, then at room temperature overnight. For each tube, 1 µl was visualized alongside a high molecular weight ladder on an agarose gel in 1x TAE buffer, and extracts showing clear bands were pooled into 4 tubes per strain. DNA extracts were cooled on ice, then digested for a second time with 21 units Promega RNase solution at 37°C for 1 hour then cooled on ice. DNA was precipitated using 0.5 volumes Promega Protein Precipitation Solution and 2 volumes of cold 96-100% ethanol on ice for 15 minutes, then pelleted at 16,162 g for 10 minutes. Pellets were washed with 3 volumes 70% ethanol, repelleted at 16,162 g for 3 minutes, and air dried for 15 minutes. DNA was resuspended in Promega DNA rehydration solution by incubation at 37°C for 1-1.5 hours. DNA extracts were spun at 16,128 g for 10 minutes, and supernatants from multiple tubes for the same strain were pooled to produce 15-30 µg genomic DNA per strain. The quality and quantity of DNA was assessed by agarose gel electrophoresis (0.8% agarose in 1x TAE buffer) alongside a high molecular weight ladder (e.g. GeneRuler High Range DNA Ladder, SM1351), by NanoDrop, Qubit fluorometer and Fragment Analyzer™ Automated CE System, and showed average DNA fragment lengths over 43 kb for NCYC 4144, and over 34 kb for NCYC 4145 and NCYC 4146.

### Long-read genome sequencing

Long-read genome sequencing was performed on the PacBio Sequel sequencing platform by the Georgia Genomics and Bioinformatics Core facility at UGA. A large insert library was constructed for each strain using the SMRTbell™ Template Prep Kit following the PacBio’s instructions for >20 kb Template Preparation using BluePippin™ Size-Selection System for Sequel™ Systems. Fragment Analyzer™ analyses showed the final size of the insert libraries was 37 kb for NCYC 4144, and 22-24 kb for NCYC 4145 and NCYC 4146. Primers were annealed to the templates, then templates were bound to polymerase using the Sequel® Binding Kit 2.0. The resultant polymerase bound complexes were purified using SMRTbell™ Clean Up Columns for Sequel™, and were then bound to MagBeads for loading. Genomic DNA for each strain was loaded onto a single SMRTcell v1 and run on the PacBio Sequel System with a movie time of 10 hours.

### *De novo* genome assembly and scaffolding

We performed *de novo* assembly with FALCON (v1.2.3) and phasing with FALCON-Unzip (v1.1.3; Chin *et al.* 2016). The assembly/phasing/polishing software were downloaded as part of the pb-assembly metapackage (v0.0.1) from bioconda (https://github.com/PacificBiosciences/pb-assembly). FALCON assembly was performed with a genome size parameter of 14.2 Mb, seed coverage of 30x, and a length cutoff for corrected reads of 1,000 bp. Consensus sequences were obtained after phasing using the Arrow polishing algorithm (v2.3.2) in FALCON-Unzip. Minimum coverage required for Arrow polishing was set to 5 and the positions of regions that could not be polished were summarized using a script (fastaLC2n.pl; https://github.com/bensassonlab/scripts) and are provided in Files S1 to S6. The full assembly pipeline was run on a computer node with 48 AMD Opteron processors and 512 GB of RAM. Software versions and parameters used for assembly are provided in Files S7 to S9.

Reference-based scaffolding of primary contigs was performed using RaGOO (v1.01; Alonge *et al.* 2019) aligned to the SC5314 reference genome (Assembly 22, A haplotype, GCF_000182965.3). RaGOO scaffolding was only used to assign primary contigs to chromosomes, and subsequent QC analysis was performed in the raw, non-scaffolded assemblies. Haplotig placement relative to the primary assembly of each strain was done by aligning haplotigs to primary contigs with minimap2 (v2.16-r922; Li 2018) using the ‘-x asm5’ alignment preset. The longest alignment for each haplotig was used to determine the placement in the primary assembly, allowing unaligned sequence from either end of the haplotig to be reported as alignment “tails”.

### Diploid assembly quality assessment

To assess their overall quality, we compared the resulting assemblies to the reference genome for *C. albicans* strain SC5314 (Assembly 22, A haplotype, GCF_000182965.3) and collected statistics using QUAST v5.0.2 (https://github.com/ablab/quast, commit 67a1136, Gurevich *et al.* 2013). Completeness of the assemblies was assessed by searching for highly conserved single-copy orthologs using BUSCO (v3.0.2, Simão *et al.* 2015) with the *Saccharomycetales* ortholog gene set from OrthoDB v9 (Zdobnov *et al.* 2016).

To obtain a visual summary of long-range assembly contiguity and correctness we performed whole genome alignments between each diploid assembly and the reference genome using minimap2 (v2.16-r922; Li 2018) with the assembly alignment preset ‘-x asm10’, and visualized them with dotPlotly (https://github.com/tpoorten/dotPlotly/). Only alignments at least 20 kb in length and with a mapping quality of 60 were visualized. For detailed analysis of structural variants in chromosome 3, we performed alignments of the diploid assemblies to the reference genome using nucmer from the MUMmer package (v4.0.0beta2, Marçais *et al.* 2018) with the settings: –maxmatch -l 80 -c 100, then visualized nucmer alignments of at least 1,000 bp using dotPlotly.

We assessed the phasing status across the genome by aligning contigs from diploid assemblies to the haploid reference and analyzing the depth of contig coverage. Alignments were obtained as described above, converted to BED format, and the depth of contig coverage along the genome was calculated using BEDTools genomecov (v2.28.0; Quinlan and Hall 2010). A visual summary of depth of contig coverage across the genome was obtained with the kpPlotCoverage function from the karyoploteR package (v1.10.2; Gel and Serra 2017). For regions of the genome where phasing was successful, we expect a depth of contig coverage of two, whereas for regions where the haplotypes were not phased, we expect a depth of coverage of one. Since phasing by FALCON-Unzip can only be done in regions were heterozygous variants are present, we also contrasted the extent of unphased regions in our assemblies (depth of contig coverage = 1) to the LOH regions detected by Bensasson *et al.* (2019) based on Illumina short-read sequencing.

For independent confirmation of phased base calls and LOH regions in each strain, we mapped Illumina HiSeq short-read data (PRJEB27862, Bensasson *et al.* 2019) to primary assemblies using bwa mem (v0.7.17; Li and Durbin 2009), produced a sorted bam file using SAMtools (v1.9; Li *et al.* 2009), then called variants using bcftools mpileup (-d 10,000) and bcftools call (-c) (v1.9; Li 2011). Because of the very high depth of Illumina sequencing for NCYC 4146, we only used data from one run (ERR2708456) for variant calling in this strain. We used vcf2allelePlot.pl (Bensasson *et al.* 2019) running R (version 3.5.0) to generate allele frequency plots for high quality base calls (phred-scaled quality >40) in order to annotate LOH regions, identify regions within the primary assemblies that had unexpectedly low heterozygosity (below 0.1% using 100 kb non-overlapping sliding windows), and summarize levels of genome-wide heterozygosity for each strain. Sites were counted as heterozygous if allele ratios were 0.5; more specifically between 0.2 and 0.8.

### Data Availability

Raw PacBio reads for the three *Candida* strains are available at the NCBI short-read archive (SRA) under BioProject PRJNA533645. The phased diploid assemblies for the three oak strains are associated with the overall BioProject PRJNA543321. Individual GenBank accession numbers for primary contigs and haplotigs, respectively, for each strain are as follows: NCYC 4144: GCA_005890765.1 and GCA_005890695.1; NCYC 4145: GCA_005890775.1 and GCA_005890685.1; NCYC 4146: GCA_005890745.1 and GCA_005890705.1. Coordinates of haplotigs relative to their respective primary assembly are available in Files S10, S11, and S12. Annotations of positions of confirmed LOH regions, assembly gaps, uncertain regions with unexpectedly low heterozygosity, and regions that were not polished by FALCON-Unzip are available for primary assemblies in Files S1, S2, and S3. Annotations of unpolished regions for alternative haplotig assemblies are provided in Files S4, S5 and S6. A full description of software version numbers for phased assembly is provided in File S7 and the configuration files used to run FALCON and FALCON-Unzip are provided in Files S8 and S9.

## RESULTS AND DISCUSSION

### High coverage long-read datasets for *C. albicans* oak strains

We generated PacBio DNA sequencing data for three strains of *C. albicans* isolated from oak trees. Each sample was independently sequenced in a single SMRT cell on the PacBio Sequel instrument. A total of 703,252, 689,752, and 863,237 subreads with average lengths of 9,082, 8,899, and 9,576 bp were obtained for strains NCYC 4144, NCYC 4145 and NCYC 4146, respectively. Theoretical coverage ranged from 415 to 558x assuming haploid coverage for a 14.8 Mb genome size (Table 1; van het Hoog *et al.* 2007). For all strains, half of the data was present in reads of approximately 16 kb or longer (Figure 1) with the longest reads obtained being 69 kb (NCYC 4144), 68 kb (NCYC 4145), and 101 kb (NCYC 4146; Table 1).

**Table 1.** PacBio sequencing statistics for *C. albicans* oak strains.

|  | NCYC 4144 | NCYC 4145 | NCYC 4146 |
| --- | --- | --- | --- |
| Yield (Gbp) | 6.4 | 6.1 | 8.3 |
| Read number | 703,252 | 689,752 | 863,237 |
| Theoretical coverage<sup>a</sup> | 432x | 415x | 558x |
| Read N50 (bp) | 16,366 | 15,964 | 16,110 |
| Longest read (bp) | 68,892 | 67,764 | 100,529 |

<sup>a</sup> Assuming a 14.8 Mb haploid genome size (van het Hoog *et al.* 2007).

**Figure 1.**
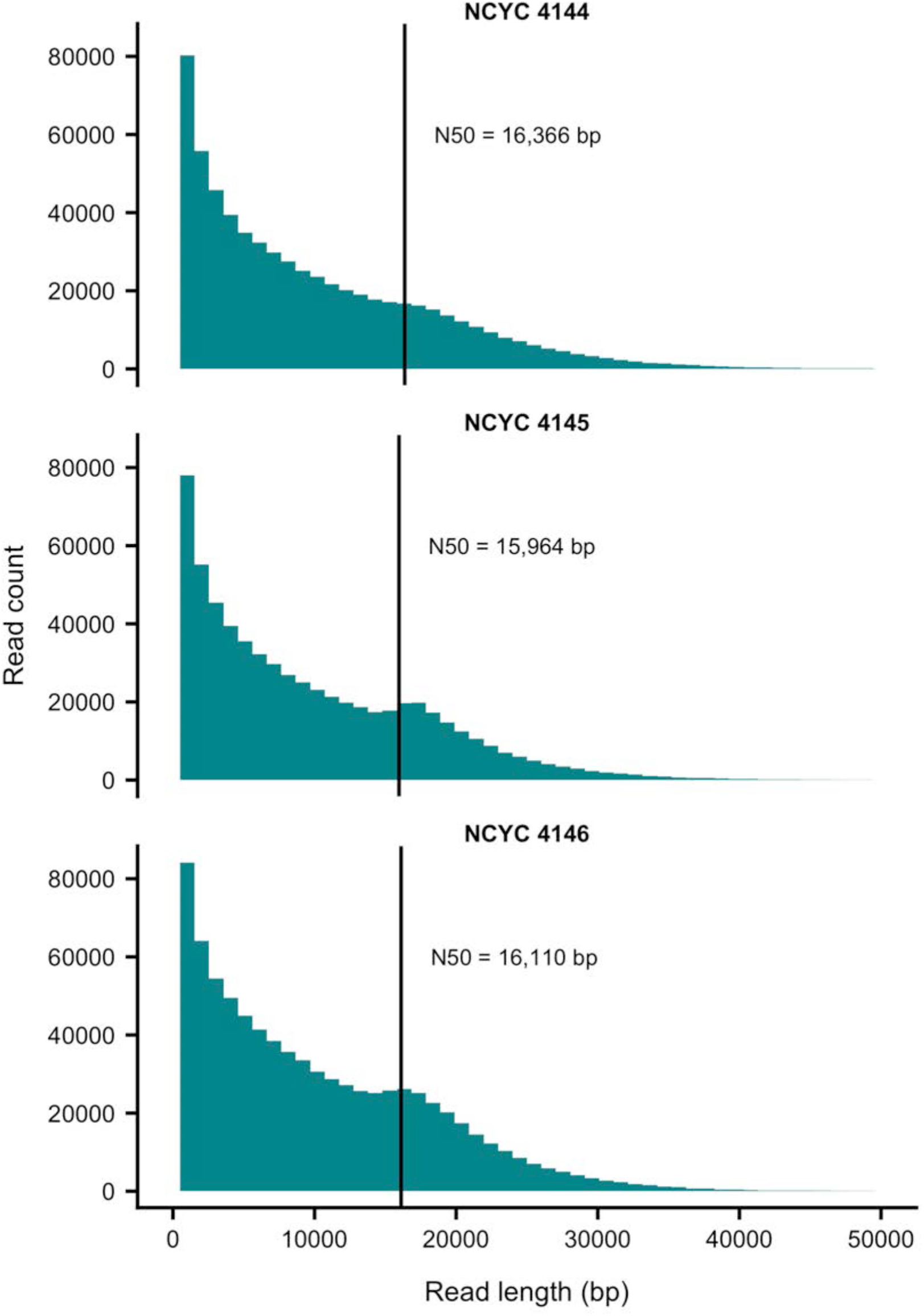
Read length distribution for the PacBio data sets generated for three *C. albicans* oak strains. Vertical lines indicate read N50 length.

### Long reads allow for high quality diploid assemblies of *C. albicans*

We used the FALCON/FALCON-Unzip assembly and phasing pipeline (Chin *et al.* 2016) to obtain phased diploid assemblies for the three *C. albicans* oak strains. For each strain, the assembly is composed of two sets of contigs; a primary contigs set, and an alternate haplotigs set. The primary contigs provide a pseudo-haploid representation of the genome, (i.e., they may include haplotype switches), and the haplotigs represent alternate haplotypes in the regions for which phasing was achieved. Phasing interruption and haplotype switches may occur when FALCON-Unzip is no longer able to establish linkage between variants (e.g. in LOH regions). We note that FALCON-Unzip removes non-linear contigs with high copy number, and as a consequence there is no mtDNA represented in the assemblies reported here.

Primary assembly size varied from 14.7 Mb in NCYC 4144 to 15.5 Mb in NCYC 4145 and NCYC 4146 (Table 2). The current reference assembly for strain SC5314 spans ~14.2 Mb, whereas the predicted genome size for *C. albicans* obtained from physical and optical maps is ~14.8-14.9 Mb (Jones *et al.* 2004; van het Hoog *et al.* 2007). Thus, the primary assembly for NCYC 4144 is very close to the predicted haploid reference genome size for *C. albicans*, while NCYC 4145 and NCYC 4146 have an additional ~500 kb of sequence in their primary assemblies. Haplotig assembly size varied from 12.5-13.7 Mb (Table 2), indicating that the vast majority (84%-92%) of the genome was phased in each assembly.

**Table 2.** *De novo* genome assembly statistics.

|  | NCYC 4144 |  | NCYC 4145 |  | NCYC 4146 |  |
| --- | --- | --- | --- | --- | --- | --- |
|  | Primary | Haplotigs | Primary | Haplotigs | Primary | Haplotigs |
| Length (bp) | 14,699,875 | 12,507,746 | 15,453,953 | 13,773,744 | 15,483,668 | 13,063,188 |
| Count | 9 | 68 | 17 | 101 | 18 | 84 |
| GC (%) | 33.6 | 33.51 | 33.68 | 33.49 | 33.59 | 33.55 |
| N50 (bp) | 1,942,916 | 254,098 | 1,652,457 | 174,215 | 1,284,452 | 248,397 |

All primary assemblies achieved high contiguity, with primary contig N50 ranging from 1.2 Mb in NCYC 4146, to 1.6 Mb in NCYC 4145, and 1.9 Mb in NCYC 4144 (Table 2). Several chromosomes were assembled as contiguous gapless sequences spanning the entire chromosome, reflecting the low fragmentation of these assemblies. For strain NCYC 4144, chromosomes 1 through 7 were recovered as single contigs, while chromosome R was split in two contigs (Figure 2A). In strain NCYC 4145, chromosomes 1, 3, 4, and 6 were recovered as single contigs whereas the remaining chromosomes were split into either 3, 5, or 9 contigs (Figure 2B). In strain NCYC 4146, chromosomes 3, 4, and 5 were recovered as single contigs whereas the remaining chromosomes were split into either 2, 4, or 5 contigs (Figure 2C). Haplotig N50 (the N50 length of phased blocks) ranged from 174 kb in NCYC 4145 to 254 kb in NCYC 4144 (Table 2). Although FALCON-Unzip produces many haplotigs per chromosome instead of fully phased chromosomes, the phase block length achieved in these assemblies is more than two orders of magnitude larger than the average gene length in *C. albicans* (1,439 bp, Braun *et al.* 2005).

**Figure 2.**
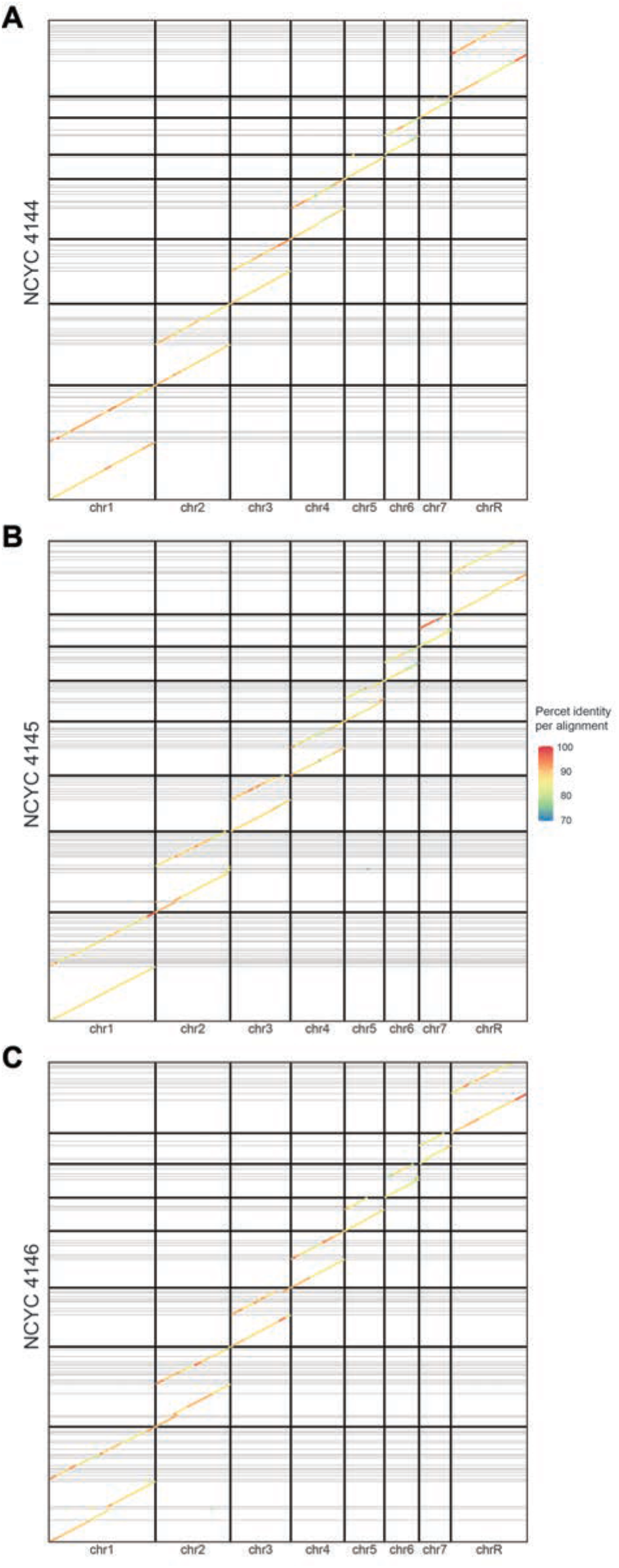
Diploid assemblies of *C. albicans* (y-axis) aligned to the reference strain SC5314 (x-axis). Darker grid lines demarcate the chromosome limits. Horizontal lines set the limits between contigs. For each chromosome, primary contigs are ordered before haplotigs and colored by percent identity to SC5314.

Whole-genome alignments and dot plots indicate large-scale genome colinearity between the reference strain SC5314 and both primary contigs or haplotigs for the oak strains (Figure 2), suggesting no major missassemblies in our assemblies or the SC5314 reference genome. BUSCO analysis on the primary contig sets revealed the presence of 97.1%, 96.2%, and 96.9% complete BUSCOs for NCYC 4144, NCYC 4145, and NCYC 4146, respectively (Table 3). We observed higher levels of completely duplicated BUSCOs in the primary assemblies for NCYC 4145 and NCYC 4146 relative to NCYC 4144, which suggests that the extra sequence in primary assemblies for these strains may be associated with duplicated segments of the genome. BUSCO analysis on the haplotig sets revealed that 79.2%, 88.2%, and 84.3% of BUSCOs are complete in the alternative haplotype of NCYC 4144, NCYC 4145, and NCYC 4146, respectively. These results indicate that our primary assemblies are highly complete in terms of gene content, and that both alleles for the majority of single copy genes are completely phased in our assemblies.

**Table 3.** BUSCO scores for the primary contigs of *C. albicans de novo* assemblies. Values represent percentages and numbers inside parentheses indicate absolute number of genes for each category.

| BUSCO category<sup>a</sup> | NCYC 4144 |  | NCYC 4145 |  | NCYC 4146 |  |
| --- | --- | --- | --- | --- | --- | --- |
|  | Primary | Haplotigs | Primary | Haplotigs | Primary | Haplotigs |
| Complete | 97.1 (1660) | 79.2 (1355) | 96.2 (1647) | 88.2 (1510) | 96.9 (1659) | 84.3 (1442) |
| Single-copy | 96.6 (1652) | 78.8 (1348) | 92.3 (1580) | 87.8 (1503) | 93.3 (1597) | 83.9 (1435) |
| Duplicated | 0.5 (8) | 0.4 (7) | 3.9 (67) | 0.4 (7) | 3.6 (62) | 0.4 (7) |
| Fragmented | 1.3 (23) | 2.1 (36) | 1.5 (26) | 2.6 (44) | 1.0 (17) | 2.0(35) |
| Missing | 1.6 (28) | 18.7 (320) | 2.3 (38) | 9.2 (157) | 2.1 (35) | 13.7 (234) |

<sup>a</sup> The *Saccharomycetales* ortholog gene set v9 was used in this analysis and comprises a total of 1,711 genes.

We verified the quality of base calls in primary assemblies by mapping independently-generated Illumina sequence (Bensasson *et al.* 2019) to the corresponding primary PacBio assembly for each strain. This resulted in high quality Illumina base calls for over 14 million invariant nucleotide sites in each strain where base calls from Illumina data matched the primary PacBio assembly. Furthermore, we observed over 70,000 sites with high quality SNP variants for each strain, which almost exclusively had the allele ratio of ~0.5 that is expected when mapping a heterozygous diploid strain to one of its parental haplotypes. All strains had fewer than 270 high quality variants in Illumina data with allele ratios of >0.95; we expect such allele ratios of ~1 if the base call in the primary PacBio assembly is incorrect. The rate of these probable errors is lower than one every 50 kb or one every 300 SNPs and confirms that the base quality of the majority of sites in the assemblies is high. For the strain with the most complete primary assembly (NCYC 4144), an unexpectedly large proportion of these errors (66 out of 169 errors; 39%) occur in the repetitive fraction of the primary assembly (21%) that is presented in lower case in the NCBI assemblies (Binomial exact test, p-value = 1 × 10^−7^). In the other two strains, a large proportion (>29%) of these errors (allele ratio ~1) occur in the fraction of the primary assembly (1%) that could not be polished using FALCON-Unzip (Binomial exact test, p-value < 2 × 10^−16^). The positions of the unpolished regions in all primary and alternate assemblies are provided in Files S1 to S6.

Heterozygous SNPs identified by mapping Illumina reads to primary PacBio assemblies also allowed us to confirm previously-identified LOH regions and estimates of genome-wide heterozygosity for each strain (Bensasson *et al.* 2019). However, patterns of SNP variation also revealed that the primary assemblies for strains NCYC 4145 and NCYC 4146 contained six regions each that showed unexpectedly low heterozygosity at (or near) breaks in the assembly not previously identified as LOH regions in Bensasson *et al.* (2019). These “uncertain” regions could represent poorly resolved structural variants or assembly errors and may explain the excess sequence and increased levels of duplication observed in NCYC 4145 and NCYC 4146. Uncertain regions span ~1 Mb in the primary assemblies of NCYC 4145 and NCYC 4146 and are described more fully in the annotations for primary assemblies along with the positions of confirmed LOH regions, gaps and unpolished regions (Files S1-S3).

### Phasing efficiency, loss of heterozygosity and aneuploidy

Whole genome alignments indicate that some regions of the genome for each strain are not phased, such as chromosome 5 in strain NCYC 4144 (Figure 2A). The phasing efficiency of FALCON-Unzip depends on the presence of heterozygous variants captured by long reads. Accordingly, variants separated by long tracts of homozygosity cannot be phased by this approach (Chin *et al.* 2016). Since the strains analyzed here were previously shown to contain segmental and whole-chromosome loss of heterozygosity (Bensasson *et al.* 2019), FALCON-Unzip is not expected to phase the entirety of these genomes.

Approximately 16.1%, 4.9% and 7.7% of the genomes of NCYC 4144, NCYC 4145, and NCYC 4146, respectively, were defined as having LOH based on variants detected in short read data relative to the SC5314 reference genome (Bensasson *et al.* 2019). As expected if LOH regions impact FALCON-Unzip phasing, the strain with the highest amount of LOH would have the lowest complete BUSCO score for haplotigs, and *vice versa*, which is demonstrated with NCYC 4144 compared to NCYC 4145 and NCYC 4146 (Table 3). To more directly analyze the correspondence between the LOH regions defined previously and the phasing efficiency of FALCON-Unzip, we mapped all contigs (primary and haplotigs) from diploid assemblies to the haploid SC5314 reference genome and analyzed the resulting depth of contig coverage profiles. While the average depth of contig coverage is approximately 2 in the non-LOH regions for the three strains, this value drops to an average of 1.31, 1.58, and 1.39 in LOH regions of NCYC 4144, NCYC 4145, and NCYC 4146, respectively (Figure 3). The difference in average depth of coverage between LOH and non-LOH regions is significant at a 99% confidence level (Welch two sample t-test, p-value < 0.001). These results confirm the expectation that homozygous regions disrupt phasing by FALCON-Unzip.

**Figure 3.**
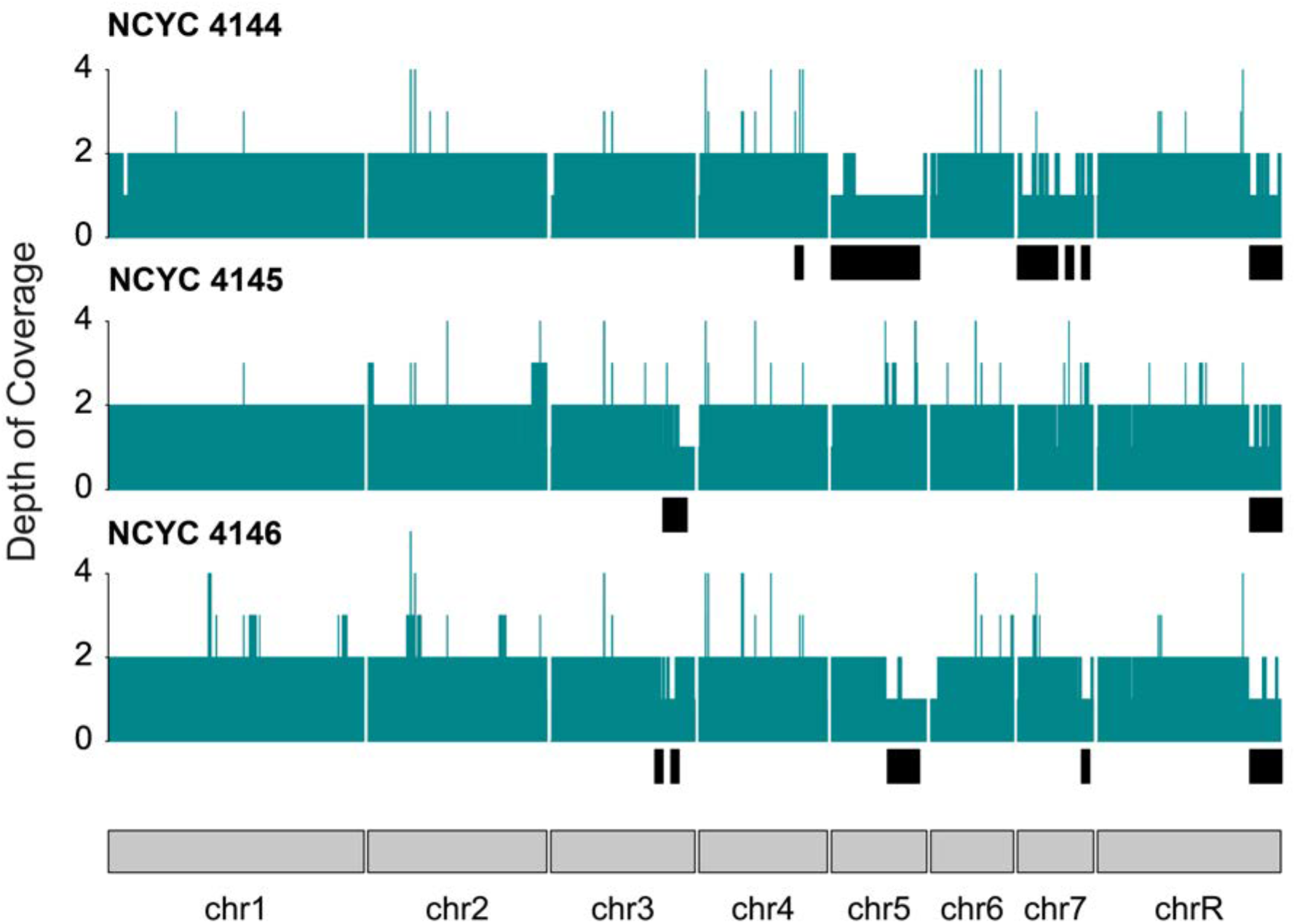
Depth of contig coverage for three diploid *C. albicans* assemblies mapped to the haploid reference genome (strain SC5314). Black boxes below each coverage plot highlight the regions defined as loss of heterozygosity in Bensasson *et al.* (2019) and generally correspond to regions of contig depth of coverage = 1.

Although largely matching the previously known LOH regions, contig coverage profiles sometimes reveal windows for which the observed depth is >2, or where the depth is not as expected given the presence or absence of LOH (Figure 3). Since contig coverage is calculated after aligning the *de novo* assemblies to the haploid reference, duplications in either haplotype that align to the same location in the reference genome could result in contig depth >2 (see subsection below). Additionally, regions defined as LOH by short-read variants that have the expected diploid contig coverage could represent regions where haplotype divergence at the nucleotide level is low, but more substantial at the structural level. Indeed, FALCON-Unzip also uses heterozygous structural variants to phase assemblies, which could potentially be used even in the absence of significant nucleotide divergence (Chin *et al.* 2016).

Allele ratio analyses in Bensasson *et al.* (2019) revealed that chrR is likely trisomic in strains NCYC 4144 and NCYC 4145. In our assemblies, the majority of chrR could be assembled and phased in two haplotypes (Figure 2 and 3). Together, these results suggest that chrR trisomy is likely represented by two major haplotypes, one with a ploidy of 2 and the other with a ploidy of 1.

### Phased diploid assemblies reveal structural variants

The evolution of *C. albicans* is characterized by frequent genomic rearrangements often involving open reading frames (ORFs) that are flanked by inverted repeats (Todd *et al.* 2019). For example, there are two groups of ORFs in neighboring regions of chromosome 3 which appear to have undergone duplication followed by inversion, generating inverted repeats (van het Hoog *et al.* 2007; Todd *et al.* 2019), and these can lead to further large-scale rearrangements (Todd *et al.* 2019). Close inspection of whole-genome alignments between the oak strains’ diploid assemblies relative to the *C. albicans* reference genome in this region reveals that oak strains have experienced inversions of the sequence between these inverted repeats (Figure 4A). The inversions are polymorphic among the three strains, with each strain displaying a unique genotype. Strains NCYC 4146 and NCYC 4144 are heterozygous for the first and second inversions, respectively, whereas strain NCYC 4145 is homozygous for both inversions. Moreover, the presence of inverted repeats in either the standard or inverted configuration results in contig depth >2 relative to the SC5314 reference genome (Figure 4B), providing an explanation for the peaks in contig coverage >2 seen in (Figure 3).

**Figure 4.**
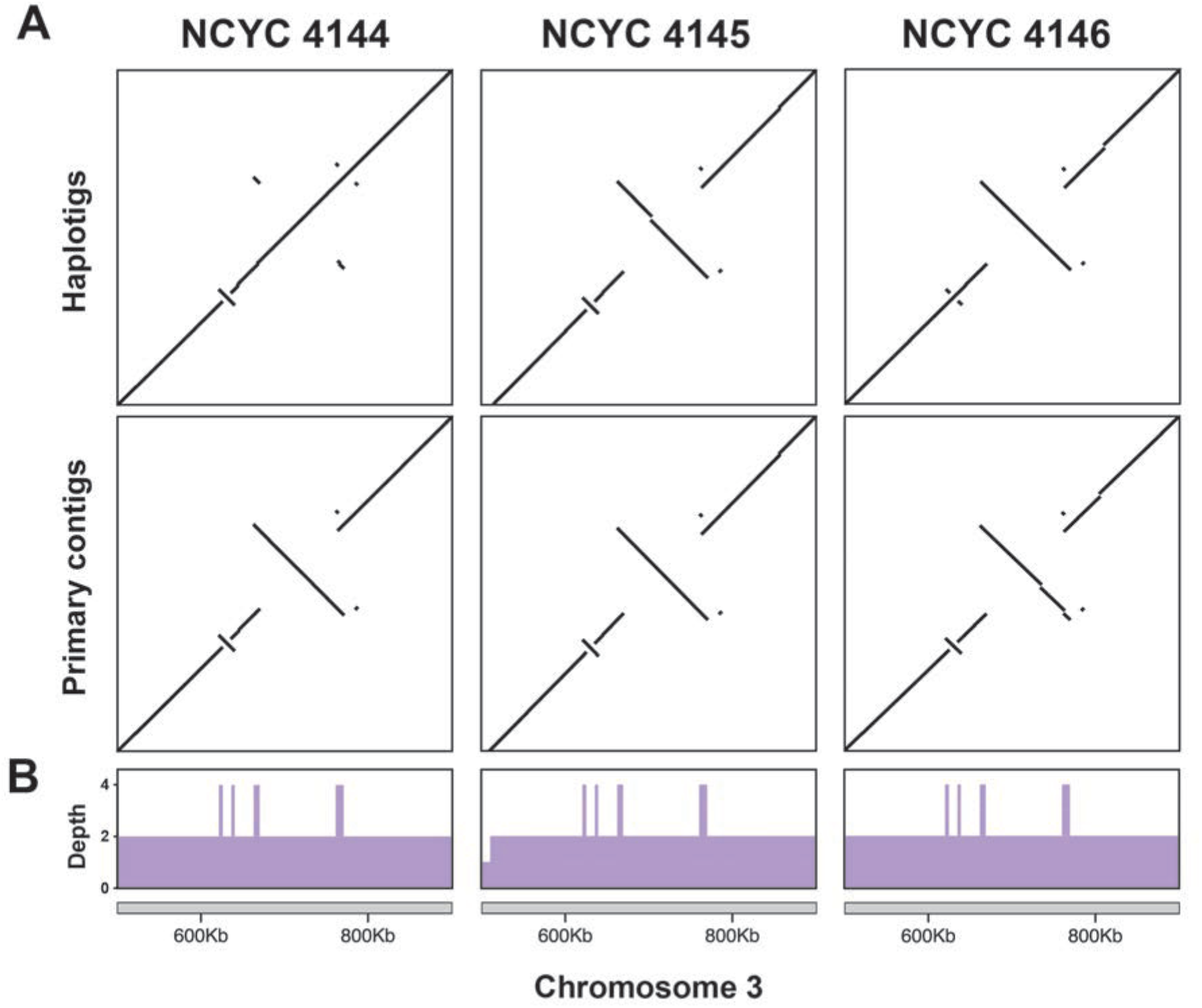
Diploid assemblies of chromosome 3 aligned to the haploid reference genome (strain SC5314). (A) Dot plots of both primary contigs and haplotigs showing a small homozygous inversion in strain NCYC 4144 followed by a large heterozygous inversion. The pattern is reversed in strain NCYC 4146, while NCYC 4145 is homozygous for both inversions. (B) Depth of contig coverage over the same region. The inverted repeats on both sides of the inverted segments display peaks with depth of coverage of 4.

## CONCLUSIONS

Here we show that PacBio SMRT sequencing is an affordable and efficient approach to obtain phased diploid assemblies for the model yeast species *C. albicans*. Our results suggest that SMRT sequencing and FALCON-Unzip assembly of the SC5314 reference strain is now merited and could provide an end to the “long hard road” to generate a complete diploid *C. albicans* reference genome (Nantel 2006). Our work also demonstrates that phased diploid assemblies generated using PacBio long-read data can provide detailed insights into genome structure and evolution in *C. albicans* that are not possible to obtain using short read sequencing. This advance is especially important for an asexual diploid species such as *C. albicans* with unusually high rates of structural rearrangements (Birky 1996; van het Hoog *et al.* 2007; Todd *et al.* 2019) that are associated with gene family evolution and pathogenicity (Butler *et al.* 2009; Todd *et al.* 2019).

## Supporting information

FileS1_NCYC4144A1_GCA_005890765.1.txt

FileS7_software.pdf

FileS8_configFalcon.txt

FileS9_configUnzip.txt

FileS10_NCYC4144placement.tsv

FileS11_NCYC4145placement.tsv

FileS12__NCYC4146placement.tsv

## ACKNOWLEDGMENTS

This study was supported in part by resources and technical expertise from the Georgia Advanced Computing Resource Center. We thank the Georgia Genomics and Bioinformatics Core, which provided the PacBio sequencing service. This work was funded by the University of Georgia.

## SUPPLEMENTAL MATERIAL

1. FileS1_NCYC4144A1_GCA_005890765.1.gff: Annotations for the NCYC 4144 primary assembly (GCA_005890765.1) in GFF format showing the positions of Loss of Heterozygosity (LOH) regions, a gap on chromosome R, and sequence that was not polished by FALCON-Unzip.
2. FileS2_NCYC4145A1_GCA_005890775.1.gff: Annotations for the NCYC 4145 primary assembly (GCA_005890775.1) in GFF format showing the positions of LOH regions, sequencing gaps, uncertain regions that show LOH but were not confirmed with independent data, and sequence that was not polished by FALCON-Unzip.
3. FileS3_NCYC4146A1_GCA_005890745.1.gff: Annotations for the NCYC 4145 primary assembly (GCA_005890745.1.gff) in GFF format showing the positions of LOH regions, sequencing gaps, uncertain regions that show LOH but were not confirmed with independent data, and sequence that was not polished by FALCON-Unzip.
4. FileS4_NCYC4144B1_GCA_005890695.1.gff: Annotations for the NCYC 4144 alternate haplotig assembly (GCA_005890695.1.gff) in GFF format showing the positions of sequence that was not polished by FALCON-Unzip.
5. FileS5_NCYC4145B1_GCA_005890685.1.gff: Annotations for the NCYC 4145 alternate haplotig assembly (GCA_005890685.1.gff) in GFF format showing the positions of sequence that was not polished by FALCON-Unzip.
6. FileS6_NCYC4146B1_GCA_005890705.1.gff: Annotations for the NCYC 4145 alternate haplotig assembly (GCA_005890685.1.gff) in GFF format showing the positions of sequence that was not polished by FALCON-Unzip.
7. FileS7_software.pdf: File in pdf format showing the software versions in the pb-assembly conda environment that were used for assembly, phasing, and polishing.
8. FileS8_configFalcon.txt: The configuration file in text format used for FALCON assembly.
9. FileS9_configUnzip.txt: The configuration file in text format used for FALCON-Unzip.
10. FileS10_NCYC4144placement.tsv: Table file in text format with tab separated values summarizing the placement of haplotigs relative to the primary assembly for NCYC 4144. Columns show the haplotig identifier created by the Arrow tool of FALCON-Unzip, chromosome number, orientation, start position, stop position, start tail, and stop tail columns. The largest single alignment was used for haplotig placement, and the length of the sequences at either side of this alignment is reported as start and stop tails.
11. FileS11_NCYC4145placement.tsv: Table file in text format with tab separated values summarizing the placement of haplotigs relative to the primary assembly for NCYC 4145. Columns show the haplotig identifier created by the Arrow tool of FALCON-Unzip, chromosome number, orientation, start position, stop position, start tail, and stop tail columns. The largest single alignment was used for haplotig placement, and the length of the sequences at either side of this alignment is reported as start and stop tails.
12. FileS12_NCYC4146placement.tsv: Table file in text format with tab separated values summarizing the placement of haplotigs relative to the primary assembly for NCYC 4146. Columns show the haplotig identifier created by the Arrow tool of FALCON-Unzip, chromosome number, orientation, start position, stop position, start tail, and stop tail columns. The largest single alignment was used for haplotig placement, and the length of the sequences at either side of this alignment is reported as start and stop tails.

