## Supplementary material for "Phased diploid genome assemblies for three strains of *Candida albicans* from oak trees": FileS7_software.pdf

| # Name | Version | Build | Channel |
|---|---|---|---|
| asn1crypto | 0.24.0 | py27_0 | |
| avro | 1.8.0 | py27_0 | bioconda |
| bcftools | 1.9 | h4da6232_0 | bioconda |
| bedtools | 2.27.1 | he941832_2 | bioconda |
| blas | 1.0 | mkl | |
| blasr | 5.3.2 | hac9d22c_3 | bioconda |
| blasr_libcpp | 5.3.1 | hac9d22c_2 | bioconda |
| bwa | 0.7.17 | ha92aebf_3 | bioconda |
| bzip2 | 1.0.6 | h14c3975_5 | |
| ca-certificates | 2018.03.07 | 0 | |
| certifi | 2018.8.24 | py27_1 | |
| cffi | 1.11.5 | py27he75722e_1 | |
| chardet | 3.0.4 | py27_1 | |
| cryptography | 2.3.1 | py27hc365091_0 | |
| curl | 7.61.0 | h84994c4_0 | |
| cython | 0.28.5 | py27hf484d3e_0 | |
| decorator | 4.3.0 | py27_0 | |
| enum34 | 1.1.6 | py27_1 | |
| falcon-kit | 1.2.3 | pypi_0 | pypi |
| falcon-phase | 1.0.0 | pypi_0 | pypi |
| falcon-unzip | 1.1.3 | pypi_0 | pypi |
| future | 0.16.0 | py27_2 | |
| genomicconsensus | 2.3.2 | py27_1 | bioconda |
| h5py | 2.8.0 | py27h989c5e5_3 | |
| hdf5 | 1.10.2 | hba1933b_1 | |
| htslib | 1.7 | 0 | bioconda |
| idna | 2.7 | py27_0 | |
| intel-openmp | 2019.0 | 118 | |
| ipaddress | 1.0.22 | py27_0 | |
| iso8601 | 0.1.12 | py27_1 | |
| libcurl | 7.61.0 | h1ad7b7a_0 | |
| libdeflate | 1.0 | h470a237_0 | bioconda |
| libedit | 3.1.20170329 | h6b74fdf_2 | |
| libffi | 3.2.1 | hd88cf55_4 | |
| libgcc | 7.2.0 | h69d50b8_2 | |
| libgcc-ng | 8.2.0 | hdf63c60_1 | |
| libgfortran-ng | 7.3.0 | hdf63c60_0 | |
| libssh2 | 1.8.0 | h9cfc8f7_4 | |
| libstdcxx-ng | 8.2.0 | hdf63c60_1 | |
| linecache2 | 1.0.0 | py27_0 | |
| minimap2 | 2.12 | ha92aebf_0 | bioconda |
| mkl | 2019.0 | 118 | |
| mkl_fft | 1.0.4 | py27h4414c95_1 | |
| mkl_random | 1.0.1 | py27h4414c95_1 | |
| mummer4 | 4.0.0beta2 | pl526hfc679d8_3 | bioconda |
| ncurses | 6.1 | hf484d3e_0 | |
| networkx | 2.1 | py27_0 | |
| nim-falcon | 0.0.0 | 0 | bioconda |
| numpy | 1.15.1 | py27h1d66e8a_0 | |
| numpy-base | 1.15.1 | py27h81de0dd_0 | |
| openssl | 1.0.2p | h14c3975_0 | |
| pb-assembly | 0.0.1 | py27_0 | bioconda |
| pb-dazzler | 0.0.0 | h470a237_0 | bioconda |

| | | | |
|---|---|---|---|
| pb-falcon | 0.2.3 | py27ha92aebf_0 | bioconda |
| pbalign | 0.3.1 | py27_0 | bioconda |
| pbbam | 0.18.0 | h1310cd9_1 | bioconda |
| pbcommand | 1.1.1 | py27h24bf2e0_1 | bioconda |
| pbcore | 1.5.1 | py27_1 | bioconda |
| perl | 5.26.2 | h14c3975_0 | |
| pip | 10.0.1 | py27_0 | |
| pycparser | 2.18 | py27_1 | |
| pyopenssl | 18.0.0 | py27_0 | |
| pysam | 0.14.1 | py27hae42fb6_1 | bioconda |
| pysocks | 1.6.8 | py27_0 | |
| python | 2.7.15 | h1571d57_0 | |
| python-consensuscore | 1.1.1 | py27h02d93b8_1 | bioconda |
| python-consensuscore2 | 3.1.0 | py27_1 | bioconda |
| python-edlib | 1.2.3 | py27h470a237_1 | bioconda |
| python-intervaltree | 2.1.0 | py_0 | bioconda |
| python-msgpack | 0.5.6 | py27h470a237_0 | bioconda |
| python-sortedcontainers | 2.0.4 | py_0 | bioconda |
| pytz | 2018.5 | py27_0 | |
| readline | 7.0 | h7b6447c_5 | |
| requests | 2.19.1 | py27_0 | |
| samtools | 1.9 | h8ee4bcc_1 | bioconda |
| setuptools | 40.2.0 | py27_0 | |
| six | 1.11.0 | py27_1 | |
| sqlite | 3.24.0 | h84994c4_0 | |
| tk | 8.6.8 | hbc83047_0 | |
| traceback2 | 1.4.0 | py27_0 | |
| unittest2 | 1.1.0 | py27_0 | |
| urllib3 | 1.23 | py27_0 | |
| wheel | 0.31.1 | py27_0 | |
| xz | 5.2.4 | h14c3975_4 | |
| zlib | 1.2.11 | ha838bed_2 | |

Software versions in the pb-assembly conda environment used for assembly, phasing, and polishing.
